# Intelligence and neuroticism in relation to depression and psychological distress: evidence of interaction using data from Generation Scotland: Scottish Family Health Study and UK Biobank

**DOI:** 10.1101/039545

**Authors:** LB Navrady, SJ Ritchie, SWY Chan, DM Kerr, MJ Adams, E Hawkins, DJ Porteous, IJ Deary, CR Gale, GD Batty, AM McIntosh

## Abstract

**Background:** Neuroticism is a risk factor for selected mental and physical illnesses and is inversely associated with intelligence. Intelligence appears to interact with neuroticism and mitigate its detrimental effects on physical health and mortality. However, the inter-relationships of neuroticism and intelligence for major depressive disorder (MDD) and psychological distress has not been well examined.

**Methods:** Associations and interactions between neuroticism and general intelligence (g) on MDD and psychological distress were examined in two population-based cohorts: Generation Scotland: Scottish Family Health Study (GS:SFHS, N=19,200) and UK Biobank (N=90,529). The Eysenck Personality Scale Short Form-Revised measured neuroticism and g was extracted from multiple cognitive ability tests in each cohort. Family structure was adjusted for in GS:SFHS.

**Results:** Neuroticism was associated with MDD and psychological distress in both samples. A significant interaction between neuroticism and g in predicting MDD status was found in UK Biobank (OR = 0.96, *p* < .01), suggesting that higher g ameliorated the adverse effects of neuroticism on the likelihood of having MDD. This interaction was not found in GS:SFHS. In both samples, higher neuroticism and lower intelligence were associated with increased psychological distress. A significant interaction was also found in both cohorts (GS:SFHS: ß = -0.05, *p* < .01; UK Biobank: ß = -0.02, *p* < .01), such that intelligence protected against the deleterious effect of neuroticism on psychological distress.

**Conclusions:** From two large cohort studies, our findings suggest intelligence acts a protective factor in mitigating the effects of neuroticism on risk for depressive illness and psychological distress.

## INTRODUCTION

Major depressive disorder (MDD) is the second-leading cause of disease burden worldwide (Ferrari *et al.*, 2013) but is not fully understood. Neuroticism is a partially-heritable personality trait describing the tendency for emotionality and sensitivity to stress (Matthews, 2009). Cross-sectional studies of clinical and non-clinical samples suggests a strong positive association between neuroticism and MDD (Chan *et al.*, 2007, Muris *et al.*, 2005, Roelofs *et al.*, 2008). Longitudinal studies find higher neuroticism prospectively associates with depression (Farmer *et al.*, 2008, Hirschfeld *et al.*, 1989, Kendler *et al.*, 2006) suggesting it is a key risk factor involved in the aetiology of MDD. This relationship has been repeatedly shown in MDD, and other disorders (Jylhä *et al.*, 2010, Van Os and Jones, 2001, Widiger and Trull, 1992). As such, neuroticism is widely considered to elevate the risk of psychopathology (Hirschfeld *et al.*, 1989).

General intelligence (*g*) is a variable created to reduce variation in performance across correlated cognitive tasks to a single indicator (Humphreys, 1979). Longitudinal studies suggest that lower cognitive ability in childhood/adolescence may confer vulnerability to psychiatric disorders (Gale *et al.*, 2009, Gale *et al.*, 2010, Maccabe, 2008), although research specifically pertaining to MDD is sparse (Gale *et al.*, 2008). Psychological distress is considerably more prevalent than MDD (Kessler and Wang, 2008) and is a defining symptom of depression (Beck *et al.*, 1979, Snaith, 1987). Low childhood intelligence (specifically in women) is associated with increased likelihood of experiencing future psychological distress (Gale *et al.*, 2009, Hatch *et al.*, 2007) which may precede the development of MDD (Gulliver *et al.*, 2012), although this is not a universal observation, particularly in studies accounting for socioeconomic status (SES).

The impacts of intelligence and neuroticism are potentially wide-ranging, including effects on physical health (Calvin *et al.*, 2011, Hakulinen *et al.*, 2015, Jokela *et al.*, 2013). Research linking neuroticism to health suggest individuals higher in neuroticism die earlier (Abas *et al.*, 2002, Roberts *et al.*, 2007, Shipley *et al.*, 2007), and are more physically ill (Hampson and Friedman, 2008, Smith and Gallo, 2001, Suls and Bunde, 2005). Although the majority of findings purport that higher neuroticism confers higher mortality risk, some studies purport no relationship (Huppert and Whittington, 1995, Iwasa *et al.*, 2008, Taga *et al.*, 2009), and some report that neuroticism provides a protective effect to physical health (Korten *et al.*, 1999, Weiss and Costa, 2005). Lower intelligence in childhood is a risk factor for poor physical health (Wraw *et al.*, 2015) and earlier death in adulthood (Deary *et al.*, 2010); a 1SD advantage in childhood is associated with a 24% reduction in mortality (Calvin *et al.*, 2011). The individual importance of both intelligence and neuroticism was supported by a longitudinal study of 4200 Vietnam War veterans (Weiss *et al.*, 2009) that found high neuroticism and low intelligence were separate risk factors for mortality, although these variables moderated the effect on the other. A significant interaction suggested that veterans high in neuroticism and low in cognitive ability were at particular risk of poor health and reduced lifespan.

While evidence from such studies suggest that IQ and neuroticism may interact to influence risk of mortality, whether similar interactions exist in regards to their influence on mental health outcomes is unknown. No investigation has examined how intelligence and neuroticism may interact in predicting and moderating each other’s associations with MDD and psychological distress; such an analysis will serve to clarify the causal mechanisms underlying MDD.

This study examined two large population-based cohorts - Generation Scotland: the Scottish Family Health Study (GS: SFHS) (Smith *et al.*, 2013a, Smith *et al.*, 2006) and UK Biobank (Allen *et al.*, 2012, Sudlow *et al.*, 2015). Due to the strong association of neuroticism with the risk of MDD found in previously (Jylhä and Isometsä, 2006), we hypothesised this effect would be replicated in our two large cohorts. Furthermore, we hypothesised that being protective against MDD, higher intelligence would confer resilience to those at risk of MDD by mitigating the adverse effects of neuroticism, akin to the interaction between intelligence and neuroticism identified for mortality (Weiss *et al.*, 2009). Finally, we posit that the effects of neuroticism and intelligence on psychological distress would mirror their effects on MDD, demonstrating similar effects on subclinical symptoms.

## MATERIALS AND METHODS

### Generation Scotland: The Scottish Family Health Study (GS:SFHS)

#### Overview

Generation Scotland: The Scottish Family Health Study (GS:SFHS) is a family-based population cohort recruited from GP practices throughout Scotland; the characteristics of the sample and protocol for recruitment have been described elsewhere (Smith *et al.*, 2013a, Smith *et al.*, 2006). Briefly, this epidemiological study collected DNA, socio-demographic and clinical data from volunteers aged 18-98 from 2006 to 2011. We analysed the 19,200 individuals from this sample that had complete data on all variables of interest. Demographic information is provided in **Table 1**.

**Table 1.** *Demographic, clinical, and cognitive characteristics of GS:SFHS and UK Biobank individuals in the current study*

|  | GS:SFHS |  |  | UK Biobank |  |  |
| --- | --- | --- | --- | --- | --- | --- |
|  | Total<br>(N = 19,200) | Control<br>(N = 16,719) | Lifetime MDD<br>(N = 2,481) | Total<br>(N = 90,529) | Control<br>(N = 60,402) | Lifetime MDD<br>(N = 30,127) |
| Age | 47.16 (14.97) | 47.23 (15.27) | 46.39 (12.89) * | 56.64 (8.13) | 57.15 (8.16) | 55.60 (7.98) * |
| Sex (% female) | 59% | 57% | 72% * | 52% | 46% | 65% * |
| Neuroticism | 3.84 (3.16) | 3.45 (2.94) | 6.45 (3.32) * | 3.46 (2.86) | 2.65 (2.43) | 5.09 (2.96) * |
| GHQ score | 15.93 (8.81) | 14.93 (7.56) | 22.70 (12.77) * |  |  |  |
| PHQ score | - | - | - | 1.36 (1.91) | 0.89 (1.33) | 2.30 (2.47) * |
| Wechsler Digit Symbol Substitution Task | 72.31 (17.09) | 72.45 (17.23) | 71.44 (16.06) * | - | - | - |
| Mill-Hill Vocabulary Test | 30.06 (4.76) | 30.05 (4.75) | 30.15 (4.84) | - | - | - |
| Weschler Logical Memory Test I & II | 31.01 (8.04) | 30.99 (8.09) | 31.02 (7.68) * | - | - | - |
| Verbal Fluency Test | 25.68 (8.10) | 25.60 (8.11) | 26.21 (8.05) * | - | - | - |
| Reaction time | - | - | - | 564.00 (119.87) | 564.70 (119.98) | 562.58 (119.66) * |
| Visual memory | - | - | - | 4.04 (3.21) | 4.04 (3.23) | 4.04 (3.17) |
| Verbal-numerical reasoning | - | - | - | 6.09 (2.14) | 6.07 (2.16) | 6.12 (2.11) * |
| SIMD | 3903.82 (1851.91) | 3957.58 (1832.28) | 3541.51 (1941.03) * | - | - | - |
| Townsend Score | - | - | - | -1.37 (2.84) | -1.47 (2.77) | -1.06 (2.94) * |

Abbreviations: GS:SFHS, Generation Scotland: the Scottish Family Health Study; MDD, Major Depressive Disorder ; GHQ, General Health Questionnaire; PHQ, Patient Health Questionnaire; SIMD, the Scottish Index of Multiple Deprivation With the exception of sex, values represent Mean (SD).
\* Significantly different from controls at $p < .05$

#### Study assessments

A screening questionnaire was administered to assess for MDD symptoms. Individuals who answered *‘Yes’* to at least one of the two screening questions *(“Have you ever seen anybody for emotional or psychiatric problems?”* and *“Was there ever a time when you, or someone else, thought you should see someone because of the way you were feeling or acting?”*) were asked to complete the Structured Clinical Interview for the Diagnostic and Statistical Manual of Mental Disorders (SCID) (First *et al.*, 1997), focusing on mood disorders. Answering *‘No’* to both screening questions resulted in assignment to the control group. Those who completed the SCID but did not meet the criteria for MDD were defined as controls. The current sample consisted of 2,481 individuals with a lifetime diagnosis of MDD (13%), and 16,719 controls (87%). At recruitment, 451 participants were currently depressed, 1,251 individuals had recorded a single episode of MDD and 1,230 individuals reported having recurrent episodes.

Intelligence was measured during recruitment interviews and the methods have been previously reported elsewhere (Smith *et al.*, 2013a, Smith *et al.*, 2006). Four cognitive tests were administered. The Wechsler Digit Symbol Substitution Task (Wechsler, 1958) measured processing speed. Verbal declarative memory was measured using one paragraph from The Weschler Logical Memory Test I & II (Wechsler, 1945). The Verbal Fluency Test measured executive function (Wechsler, 1958) using phonemic lists of C, F and L – participants recalled as many words as possible beginning with each of the three letters (60 seconds was given for each letter). The Mill-Hill Vocabulary Test (Raven, 1958) was administered as a measure of vocabulary, using the combined junior and senior synonyms. From these tests, a measure of general intelligence (*g*) was extracted as the first un-rotated principal component in a principal components analysis (Marioni *et al.*, 2014). This *g* component explained 41% of the variance in the four measures of intelligence, with loadings for processing speed, vocabulary, verbal declarative memory and executive function being 0.57, 0.68, 0.63 and 0.69 respectively.

Neuroticism was measured using the neuroticism subscale of the Eysenck Personality Questionnaire Short Form-Revised (EPQ-SF) (Eysenck, 1991). Individual answers on 12 questions with a binary response (*‘Yes/No’*), yielding a total score ranging between 0 and 12 were aggregated to produce a total neuroticism score. This scale has been concurrently validated (Gow *et al.*, 2005).

The General Health Questionnaire (GHQ-28) (Goldberg and Hillier, 1979) is a screening device for identifying psychiatric symptoms. Four subscales were designed to assess: somatic symptoms, anxiety and insomnia, social dysfunction and depression. In aggregating the scores from each sub-scale using the Likert method, a total GHQ score was obtained in which higher scores indicate increased psychological distress.

The Scottish Index of Multiple Deprivation (SIMD) (Payne and Abel, 2012) is a tool for identifying deprived areas in Scotland by combining different indicators of deprivation (income, employment, health, education, and crime) into a single index. The SIMD recognises 6,505 small areas – datazones ‐ each containing approximately 350 households. A relative ranking for each datazone is provided, from 1 (most deprived) to 6505 (least deprived).

GS: SFHS received ethical approval from the NHS Tayside Committee on Medical Research Ethics (REC Reference Number: 05/S1401/89). Written consent for the use of data was obtained from all participants.

### UK Biobank

#### Overview

UK Biobank is a population cohort of more than 500,000 individuals aged 45-69 recruited across the UK from 2006-2010. The characteristics of the cohort have been previously described (Allen *et al.*, 2012, Sudlow *et al.*, 2015). Sociodemographic, mental health and cognitive data were collected from participants, in addition to administering medical assessments and collecting blood samples. Baseline assessments were extensive (Smith *et al.*, 2013b). Demographic information on these individuals is provided in **Table 1**.

During a final wave of assessment, MDD and bipolar symptoms were assessed. Although no specific psychiatric interview was undertaken, criteria for mood disorders followed the structured diagnostic approach within the International Classification of Diseases (ICD‐ 10) (World Health Organization., 1992) and the American Psychiatric Association’s Diagnostic and Statistical Manual (DSM-IV) (American Psychiatric Association, 2003). Classification for single and recurrent episode(s) of MDD followed Smith *et al.* (2013b). Single and recurrent episode MDD were collapsed to produce a single, dichotomised variable of case/control. The final cohort consisted of 90,529 individuals for whom full phenotype data was available, of which 30,127 (33%) had a lifetime diagnosis of MDD, and 60,402 (67%) were identified as controls.

#### Study assessments

Three custom-made cognitive tests were administered in UK Biobank, completed during baseline assessment. This cognitive data has been described previously (Smith *et al.*, 2013b). Reaction time was measured using a timed test of symbol matching, similar to the card game ‘Snap’. The score for analysis was the mean response time in *ms* over 12 trials; higher reaction times indicated poorer performance. A task with thirteen logic/reasoning-type questions assessed the ability to solve verbal-numerical reasoning problems. The score was the total number of correct answers given within a two-minute period. A visuospatial memory test was administered in which six pairs of cards were presented on-screen in a random pattern before being turned face down on the screen. Participants were asked to match as many pairs as possible using the fewest attempts. The score for analysis was the number of errors made; higher scores reflect poorer cognitive function. From these tests, a measure of general intelligence (*g*) was extracted as the first un-rotated principal component (Marioni *et al.*, 2014), explaining 44% of the variance in scores. Loadings onto the *g* factor were: -0.61 (verbal-numeric reasoning), 0.57 (visual memory), and 0.55 (reaction time).

Neuroticism was assessed using 12 questions from the Eysenck Personality Questionnaire Short Form-Revised (EPQ-SF) (Eysenck, 1991). Each question has a binary response (*‘Yes/No’*) with the number of items coded *‘Yes’* being aggregated to produce a total neuroticism score, with higher scores reflecting higher levels of neuroticism.

The Patient Health Questionnaire 9 (PHQ9) (Kroencke *et al.*, 2001) incorporates DSM-IV diagnostic criteria to rate the frequency and severity of depressive symptoms from the two week period prior to completion. Participants were administered the first four questions from the PHQ-9, responding on a four-point Likert scale. Scores were aggregated to produce a total PHQ score, with higher scores denoting higher levels of psychological distress.

The Townsend Deprivation Index (Townsend, 1987) is a census-based measure of deprivation. It is a tool for identifying socioeconomic status by proxy by incorporating four aspects of deprivation; unemployment, non-car ownership, non-home ownership and household overcrowding - combining them into a single. Small geographical areas based on postcode information are allocated ‘Townsend Scores’ with higher scores representing a greater degree of deprivation within that area.

UK Biobank received ethical approval from the North West Multicentre Research Ethics Committee (REC Reference Number: 11/NW/0382). Written consent for the use of data was obtained from all participants. This study was conducted under UK Biobank application 4844 “Stratifying Resilience and Depression Longitudinally” (PI Andrew McIntosh).

#### Statistical analysis

In GS:SFHS, we used the MCMCglmm software package (Hadfield, 2010) in R Studio which employs a Markov Chain Monte Carlo (MCMC) estimator for the generalised linear mixed model to allow for binary outcomes (using the “threshold” family with a probit link function). We fitted neuroticism and *g* as fixed effects to evaluate the main effect of neuroticism and *g* as an independent possible risk factor for MDD. We fitted the interaction of neuroticism and *g* to estimate the moderating effect of g on the contribution of neuroticism to MDD. We also fitted a model to examine the interaction between neuroticism and *g* while conditioning on deprivation (SIMD). Because MDD is a binary outcome we report regression coefficients as odds ratios. Furthermore, as GS:SFHS is a family-based study, MCMCglmm allowed us to control for the relatedness of the sample by fitting familial structure as a random effect by creating an inverse relationship matrix using pedigree kinship information. In a second set of analyses, we used GHQ score as the outcome variable and treated is as normally distributed. Neuroticism and GHQ scores were scaled to have a mean of zero and a standard deviation of 1, such that the reported regression coefficients (β) are standardized, allowing for easy interpretation of the results. In extracting *g* as the first principal component from the cognitive variables, *g* was also standardized to have a mean of zero and a standard deviation of one. Age, age^2^ and sex were used as fixed effects throughout.

In UK Biobank, multiple regression analyses were conducted. We examined the main effect of neuroticism on MDD and *g* as a hypothesized independent risk factor, as well as the interaction between neuroticism and *g* on the likelihood of MDD. Finally, the interaction between neuroticism and *g* adjusting for deprivation (Townsend score) was investigated. Because MDD is a binary outcome we fitted a generalized linear regression with a logit link function and report odds ratios. A second set of analyses examined psychological distress (PHQ score) as the outcome using linear regression models. Neuroticism and GHQ scores were scaled to have a mean of zero and a standard deviation of one, such that the reported regression coefficients (betas) are standardized. Reaction time was transformed with a log transformation due to a significantly positive skew. Visual memory was transformed with a log+1 transformation because it was significantly skewed and zero-inflated. *G* was also standardized to have a mean of zero and a standard deviation of one. All regression analyses co-varied for age, age^2^ and sex.

## RESULTS

### GS:SFHS

As seen in **Table 1**, MDD cases were significantly younger, had higher psychological distress (GHQ) scores, were more likely to be female, and had higher neuroticism scores. No significant differences were found between control and MDD participants’ general intelligence; t(3243.38) = −1.39, *p =* 0.17, Cohen’s *d* = .03, although group differences were found in processing speed and executive function. A significant difference was found between control and MDD participants’ deprivation; t(3171.20) = 9.93, *p* = 2.20×10^−16^, Cohen’s *d* = .22 in which cases tended to be from less deprived areas. Output for each regression model can be found in the supplementary material provided.

### Associations of neuroticism and g with MDD status

Greater neuroticism scores were significantly associated with increased likelihood of having MDD. For every 1SD increase in neuroticism score, the odds of having MDD increased by 3.62 ([95% CIs = 3.29, 4.03], *p* = < 1.00×10^−4^). Age, sex and age^2^ were also significant predictors in this model: increasing age and being female increased the likelihood of having MDD when fitted as a fixed effects. A second model indicated there were no independent effects of *g* on the likelihood of MDD (OR = 0.96, [95% CIs = 0.90, 1.01], *p* = 0.12), although age, age^2^ and sex remained significant.

### Interaction between neuroticism and g on MDD

A third model examined the interaction between neuroticism and *g*. Neuroticism was strongly associated with the likelihood of having MDD (OR = 3.68, [95% CI = 3.33, 4.09], *p* < 1.00×10^−4^) and higher *g* was associated with a moderately increased the likelihood of having MDD (OR = 1.08, [95% CIs = 1.01, 1.14], *p =* 1.54×10^−2^). There was no significant interaction between neuroticism and *g* (OR = 1.04, [95% CI = 0.99, 1.09], p = 0.12). After entering SIMD into the model, the interaction between neuroticism and *g* remained non-significant. Less social deprivation was a significant factor in reducing the likelihood of MDD (OR = 0.86, [95% CIs = 0.82, 0.90], *p* < 1.00×10^−4^).

### Associations of neuroticism and g with psychological distress

Neuroticism scores were significantly associated with increased levels of psychological distress. For every 1SD increase in neuroticism score, GHQ score increased by 0.51 ([95% CIs = 0.50, 0.53], *p* < 1.00×10^−4^). Age, sex and age^2^ were also significant predictors whereby increasing age and being female increased the likelihood of having MDD. A second model indicated that increasing *g* was associated with significantly decreased levels of psychological distress (ß = −0.09, [95% CIs = −0.10, −0.08], *p* < 1.00×10^−4^); age, age^2^ and sex remained significant.

### Interaction between neuroticism and g on psychological distress

A third model examined the interaction between neuroticism and *g* on psychological distress. Neuroticism was strongly associated with increased psychological distress (ß = 0.50, [95% CIs = 0.49, 0.51], *p* < 1.00×10^−4^) whereas *g* was significant associated with a decreased likelihood of psychological distress (ß = −0.05, [95% CIs = −0.06, −0.04], p < 1.00×10^−4^). A significant interaction between neuroticism and *g* was found whereby higher *g* mitigated the detrimental effects of neuroticism on psychological distress (ß = −0.02, 95% [CIs = −0.03, −0.02], *p* < 1.00×10^−4^). After accounting for SIMD, the interaction between neuroticism and *g* remained significant. Lower levels of deprivation were significantly associated with lower levels of psychological distress (OR = 0.95, [95% CIs = 0.94.96], *p* < 1.00×10^−4^).

### UK Biobank

Demographic information from UK Biobank is reported in **Table 1**. MDD cases were significantly younger, had higher psychological distress (PHQ) scores, were more likely to be female, and had higher neuroticism scores than controls. Significant differences between cases’ and controls’ cognitive test scores were found in verbal-numerical reasoning (in which controls performed better) and reaction time (in which cases performed better); in addition to a significant difference in *g* (t(61357) = −2.65, p = 8.12×10^−3^, Cohen‘s *d* = .02) - general intelligence was found to be higher in MDD cases. Control participants had a significantly lower average deprivation score than did participants with MDD (t(57110) = −20.08, *p* = 2.2×10^−16^), indicating that MDD cases lived in less deprived areas than controls. Full statistical output can be found in the supplementary material.

### Associations of neuroticism and g with MDD status

We fitted a logistic regression to predict the likelihood of having MDD based on neuroticism. Greater neuroticism scores were significantly associated with increased likelihood of having MDD (see **Table 2**). For every 1SD increase in neuroticism, the odds of having MDD increased by 2.39 (95% confidence intervals = [2.35, 2.43], *p* < 2.00×10^−16^). A second model indicated that there were no main effects of *g* on likelihood of MDD (OR = 0.99, [95% CIs = 0.98, 1.01], *p* = 0.39) when fitted without Neuroticism. Age, sex, and age^2^ were significant predictors in both models whereby increasing age and being female increase the likelihood of having MDD.

### Interaction between neuroticism and g on MDD

A third model examined the interaction between neuroticism and *g*. Neuroticism was strongly associated with likelihood of having MDD (OR = 2.40, [95% CI = 2.36, 2.44], *p* < 2.00×10^−16^). Greater *g* was associated with a moderately increased likelihood of having MDD (OR = 1.05, [95% CIs = 1.04, 1.07], p = 5.29×10^−12^). There was a significant interaction between neuroticism and *g.* Intelligence was a greater risk factor for MDD at lower levels of Neuroticism (OR = 0.96, [95% CIs = 0.95, 0.98], *p =* 3.44×10^−7^), see **Table 2** and **Figure 1**. After entering Townsend score into the model, the interaction between neuroticism and *g* remained significant. Socio-economic deprivation was a significant factor in increasing the likelihood of MDD (OR = 1.04, [95% CIs = 1.04, 1.05], *p* < 2.00×10^×16^).

**Fig. 1.**
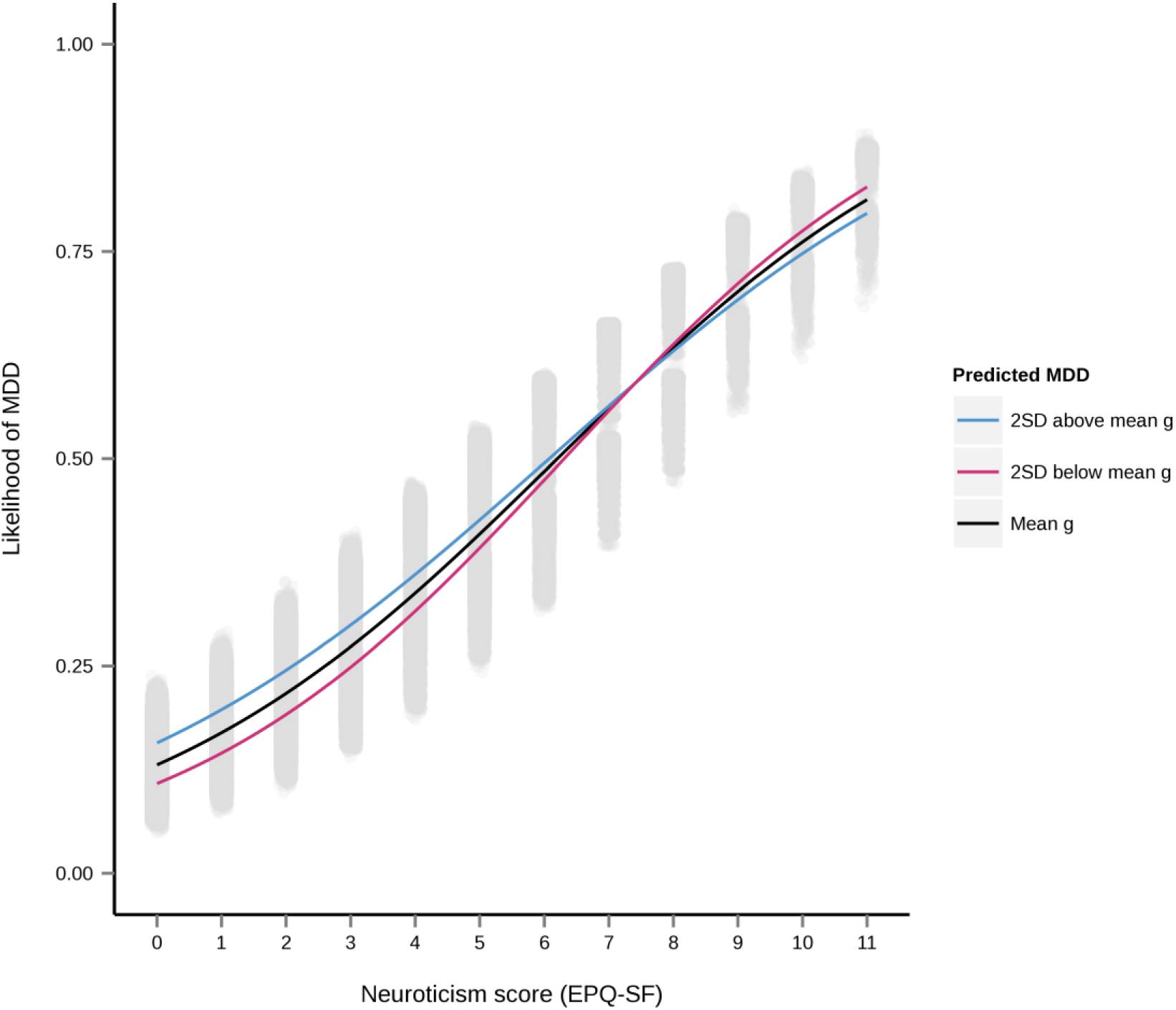
Predicted probability of having MDD from the interaction of neuroticism and *g* in UK Biobank. Regression lines reflect the interaction at mean *g* (black line) and 2SD above (blue line) and below mean *g* (pink line).

**Table 2.** *Results of a logistic regression from UK Biobank predicting odds ratios for MDD status, beta coefficients for psychological distress (PHQ), upper and lower 95% confidence intervals, the Akaike Information Criterion and adjusted R^2^ value for the model*

| Outcome | Variables | Odds Ratio | $\beta$ | Lower 95% CIs | Upper 95% CIs | $p$ value | AIC | $R^2$ |
| --- | --- | --- | --- | --- | --- | --- | --- | --- |
| MDD | Neuroticism | 2.40 *** | - | 2.36 | 2.44 | $< 2.00 \times 10^{-16}$ | 98649 | - |
| | *g* | 1.05 *** | - | 1.04 | 1.07 | $5.29 \times 10^{-12}$ | | - |
| | Age | 1.16 *** | - | 1.13 | 1.20 | $< 2.00 \times 10^{-16}$ | | - |
| | Age<sup>2</sup> | 1.00 *** | - | 1.00 | 1.00 | $< 2.00 \times 10^{-16}$ | | - |
| | Sex (M) | 0.67 *** | - | 0.65 | 0.68 | $< 2.00 \times 10^{-16}$ | | - |
| | N* *g* | 0.96 *** | - | 0.95 | 0.98 | $3.44 \times 10^{-7}$ | | - |
| PHQ | Neuroticism | - | 0.51 *** | 0.51 | 0.52 | $< 2.00 \times 10^{-16}$ | - | .2977 |
| | *g* | - | -0.05 *** | -0.06 | -0.05 | $< 2.00 \times 10^{-16}$ | | |
|  | Age | - | 0.00 | -0.00 | 0.01 | 0.55 |  |  |
| | Age<sup>2</sup> | - | -0.00 *** | -0.00 | 0.00 | $1.88 \times 10^{-4}$ | | |
| | Sex (M) | - | 0.02 *** | 0.02 | -0.00 | $6.20 \times 10^{-9}$ | | |
| | N* *g* | - | -0.02 *** | -0.03 | -0.02 | $< 2.00 \times 10^{-16}$ | | |

Abbreviations: MDD, major depressive disorder; PHQ, Patient Health Questionnaire; AIC, Akaike Information Criterion; *g*, General Intelligence
\*\*\* Significantly different from controls at $p < .001$

### Associations of neuroticism and g with psychological distress

Neuroticism scores were significantly associated with increased levels of psychological distress. For every 1SD increase in neuroticism score, psychological distress increased by 0.51 ([95% confidence intervals = 0.51, 0.52], *p* < 2.00×10^−16^). A second model indicated that increasing *g* significantly decreased levels of psychological distress (ß = -0.08, [95% CIs = -0.08, -0.07], *p* < 2.00×10^−16^). Age, age^2^, and sex remained significant across models.

### Interaction between neuroticism and g on psychological distress

A third model examined the interaction between neuroticism and *g* on psychological distress. Neuroticism was strongly associated with increased psychological distress (ß = 0.51, [95% CIs = 0.51, 0.52], *p* < 2.00×10^−16^) whilst *g* was significantly associated with decreased psychological distress (ß = −0.05, [95% CIs = −0.06, −0.05], *p* < 2e-16). A significant interaction between neuroticism and *g* was found in which *g* moderates the detrimental effects of neuroticism on psychological distress (ß = −0.02, [95% CIs = −0.03, −0.02], *p* < 2.00×10^−16^), see **Table 2** and **Figure 2**. The interaction between neuroticism and *g* remained significant after co-varying for deprivation which itself was demonstrated to be a significantly associated with increasing psychological distress (OR = 0.03, [95% CIs = 0.03, 0.03], *p* < 2.00×10^−16^).

**Fig. 2.**
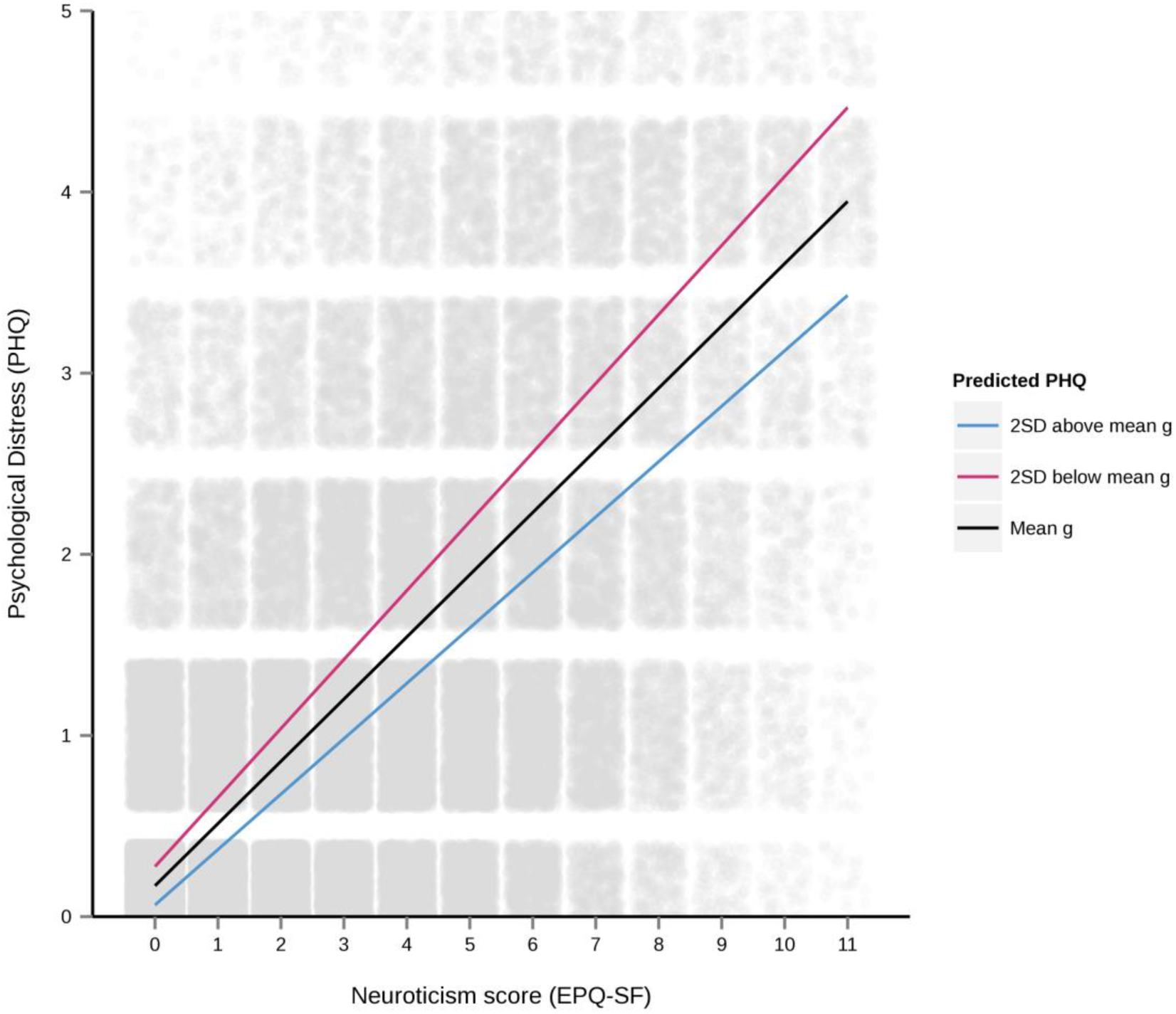
Psychological distress (PHQ) scores from the interaction of neuroticism and g in UK Biobank. Regression lines reflect the interaction at mean g (black line) and 2SD above (blue line) and below mean g (red line)

## DISCUSSION

The current study examined cross-sectional associations between neuroticism and general intelligence (*g)* and major depressive disorder (MDD) and psychological distress in two large population based cohorts; GS:SFHS and UK Biobank. Neuroticism was significantly associated with the presence of MDD, replicating previous findings (Chan *et al.*, 2007, Muris *et al.*, 2005, Ormel *et al.*, 2001). General intelligence conferred no consistent independent risk or resilience to MDD status without co-varying for neuroticism. Notably however, data from UK Biobank suggested that higher general intelligence reduced the impact of neuroticism on risk of MDD, providing resilience to those at high risk of MDD. By contrast, results from GS:SFHS suggest that intelligence does little to negate the adverse effects of neuroticism on MDD.

This discord between cohorts was not found in a second set of analyses on psychological distress. Consistent findings from both GS:SFHS and UK Biobank suggested that higher neuroticism scores were associated with increased psychological distress, whereas higher intelligence was associated with reduced levels. A significant interaction was found between neuroticism and general intelligence in both samples such that higher neuroticism was associated with lower psychological distress scores in those with greater intelligence. Our findings from 109,729 individuals across two cohorts suggest that high on neuroticism and high in general intelligence reduce susceptibility to psychological distress and that higher general intelligence lessens the strength of the neuroticism-distress association.

### Prior studies

To our knowledge, this is the first empirical association of intelligence providing a protective function to those at high risk of MDD (by virtue of high neuroticism) and psychological distress in the phenotypic domain literature. Confirming previous research (Akiskal *et al.*, 1983, Hirschfeld *et al.*, 1989, Jylhä *et al.*, 2010, Van Os and Jones, 2001, Widiger and Trull, 1992), we found that higher neuroticism is associated with increased likelihood of MDD. The inverse association found here between intelligence and MDD confirms previous longitudinal work (Gale *et al.*, 2010, Gale *et al.*, 2008, Koenen *et al.*, 2009, Maccabe, 2008). Akin to the Vietnam Experience Study (Weiss *et al.*, 2009), this study provides evidence for the association of intelligence as a protective factor in those at risk for MDD (i.e., high neuroticism).

Whereas intelligence confers an apparent resilience to those at high risk of MDD, how it does so is not clear. Intelligence could be a marker of better system integrity (Deary, 2012); increased intelligence could circumvent the biasing of mood toward negative emotion in individuals high in neuroticism that may lead to psychological distress and MDD symptomology (Hasler *et al.*, 2004). Alternatively, more intelligent individuals may be better able to employ successful coping mechanisms during times of psychological distress and illness; brighter children are found to be more resilient in the face of adversity (Fergusson *et al.*, 2005). Research also suggests that psychosocial factors (such as coping style) are associated with resilience to mood and anxiety disorders (Garmezy *et al.*, 1984, Masten and Coatsworth, 1998). More intelligent individuals may voice their distress more readily to caring professionals (task-oriented coping) that may, in turn, decrease psychological distress and future likelihood of MDD. This possibility is also consistent with the finding that whilst *g* and neuroticism interacted to reduce psychological distress, they did not interact to predict MDD in both cohorts. The influences of coping style (Cosway *et al.*, 2000, Higgins and Endler, 2006, McWilliams *et al.*, 2003) on intelligence should be examined in future investigations as it could be a mediating factor in the depressogenic process. Furthermore, individuals with higher intelligence may live in less deprived areas with better access to [mental] healthcare; an assertion supported by our findings in GS:SFHS whereby less deprivation is associated with a reduction in both MDD and psychological distress.

Increased psychological distress is an established symptom of depression and is often used as the basis of diagnosis (Beck *et al.*, 1979, Snaith, 1987). More intelligent individuals may be better able to decrease the intensity of their psychological distress having established effective coping strategies during previous exposure (reviewed in (LeDoux and Gorman, 2001). Goldberg *et al.*, (1987) described distress as representing the overall severity of MDD and so with increased psychological distress, the probability of having MDD may also increase.

### Study strengths and limitations

Some possible caveats to this study merit comment. There is currently a partial disparity in cognitive tasks used to generate *g* in our samples. GF:SFHS used pre-existing, standardized measures whereas UK Biobank generated bespoke cognitive tasks, novel to that particular cohort. The findings from this study would benefit from a meta-analysis, to examine the similarity of results across samples.

A second limitation is that the MDD phenotype was defined differently across samples. GS:SFHS determined MDD diagnosis using SCID (First *et al.*, 1997), and as such a robust MDD phenotype was obtained. However, UK Biobank aggregated self-reported depression questions to form a depression phenotype; this data is not as comprehensive. The disparity between MDD phenotypes and the familial structure fitted in GS:SFHS may explain the imbalance in results found in this investigation. The association of intelligence in reducing MDD likelihood requires further exploration and validation.

Additionally, this investigation only examined the personality trait neuroticism. Personality represents stable individual dispositions in emotional reactivity, behavioural tendencies, and cognitive styles (Deary *et al.*, 2010, Roberts *et al.*, 2007), all of which may be mediated/moderated by intelligence in mental health. Thus, examining the associations between all major dimensions of personality, general intelligence and MDD in subsequent research is advised.

In conclusion, two large population-based cohorts demonstrated that intelligence associates as a protective factor in mitigating the effects of neuroticism on risk of depressive illness and psychological distress. Future research should determine the relationship prospectively in a sample where incident cases of MDD can be identified. An important corollary of this work may inform the stratification of depression, and future studies designed to disentangle the cognitive mechanisms driving depression are an important next step in further elucidating the aetiology of the disorder.

## ACKNOWLEDGEMENTS

This investigation was supported by Wellcome Trust Grant 104036/Z/14/Z and by the Dr Mortimer and Theresa Sackler Foundation. This research has been conducted using the UK Biobank Resource established using funding from the Wellcome Trust, Medical Research Council, Scottish Government and others. Generation Scotland received core support from the Chief Scientist Office of the Scottish Government Health Directorates [CZD/16/6] and the Scottish Funding Council [HR03006].

